# ybyra: Y-chromosome haplogroup calling using a tree-based scoring method

**DOI:** 10.1101/2025.11.20.689455

**Authors:** Thomaz Pinotti, Hugh McColl, Martin Sikora, Lucas Czech

## Abstract

We present *ybyra*, a Snakemake workflow for automated Y-chromosome haplogroup assignment using a tree-based scoring method with robust and transparent heuristics. The pipeline supports both human reference genome builds 37 and 38, makes use of three different well-curated Y-SNP tree topologies and features an optional ancient DNA damage filter to accommodate low-coverage degraded samples. *ybyra* produces reproducible, scalable haplogroup calls with detailed scoring outputs to facilitate quality assessment and reporting. We further showcase its versatility by applying it to horse Y-chromosome data, highlighting its applicability beyond humans.

**Availability:** ybyra is published under the MIT license and freely available at github.com/tpinotti/ybyra.

## 1. Introduction

The Y-chromosome is the largest uniparentally inherited haplotypic block in the genome of mammals^1,2^ and has been widely used in phylogenetics, forensics and in reconstructing population history in humans and other species^3–8^. With the rapid development of the human Y-chromosome phylogenetic tree, it has been become standard – often required – in population genetics and ancient DNA studies to report the specific node of the human Y-chromosome tree (known as *haplogroup*^9,10^) to which each individual belongs. Identifying an individual’s haplogroup allows for its placement within this tree and provides insight into lineage relationships, migration and demographic history^5^ as well as allowing for a more robust pedigree reconstruction^11,12^.

Accurate haplogroup identification depends on reliable SNP calling, integration of both derived and ancestral calls and placement into a well-curated Y-chromosome phylogenetic tree. Existing tools are often limited by rigid tree structures, lack support for multiple topologies or are not optimized for ancient DNA (aDNA) datasets characterized by low-coverage and post-mortem modifications^13–16^. They also do not produce interpretable outputs for quality control and reproducible reporting.

To address these limitations, we developed *ybyra*, a reproducible workflow for Y-haplogroup inference that by default 1) supports both GRCh37 and GRCh38 builds, 2) accommodates three widely used and well-curated Y-SNP tree topologies and 3) implements an optional aDNA damage filter. It can also be easily customized for other tree topologies, including non-human species.

## 2. Pipeline overview

### 2.1. Input

*ybyra* is implemented as a Snakemake workflow^17^, as shown in **Figure 1**, enabling scalable execution on local or cluster computing environments. The input consists of BAM files^18^ aligned to either GRCh37 or GRCh38 (or any custom reference, see below). Parameters – such as reference build, tree choice and ancient DNA damage filtering – are specified through a YAML configuration file.

**Figure 1:**
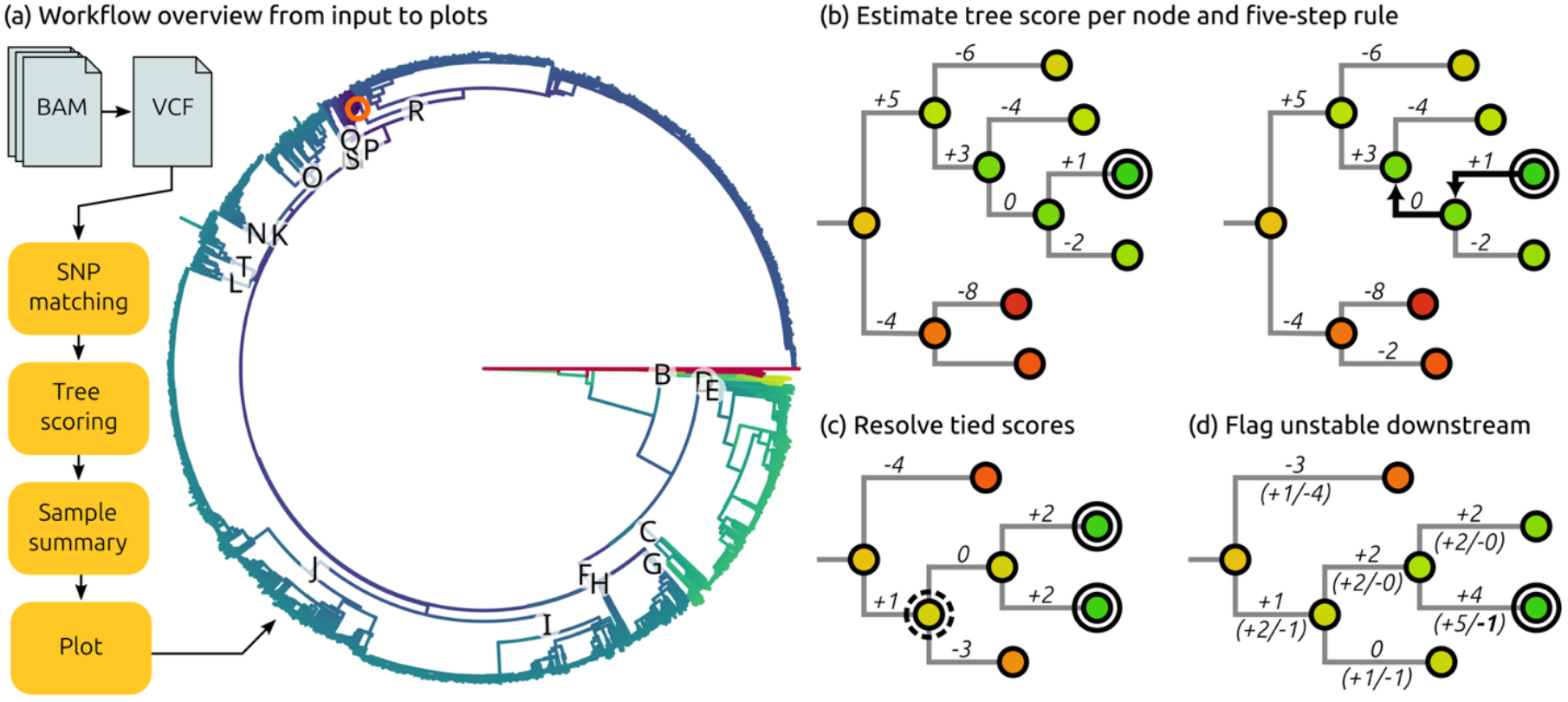
Overview and scoring process. (a) *ybyra* takes BAM files of individual samples as input and uses BCFtools to call and query SNPs. These are then matched with the tree (optionally assessing damage), and a score is computed for every node in the tree (optionally filtering damaged calls). The per-node scores are then summarized into a best placement per sample and plotted. We offer detailed output and plots for manual inspection, as well publication-ready visualizations. Here, we show the human tree annotated with major haplogroups and colored by the per-node score, and the highest scoring node for the sample in the orange circle. (b) Details of the sample summary step: For a sample, the tree score is calculated for all nodes by aggregating the derived (+1) or ancestral (-1) calls along the branches. The highest scoring node (encircled) is taken as the placement of the sample. We then apply a heuristic “five-step rule”, checking if the placement is supported by another derived SNP at most five steps upstream. Here, this is the case within two steps (+3 score at the second node upstream). (c-d) Special cases. (c) In case of a tie, we choose the most recent common ancestor with a derived hit as the best placement (dashed circle) and flag the sample accordingly. (d) The derived and ancestral scores contributing to the aggregation are shown here in detail per node. If there is an ancestral call at the highest-scoring node (bold -1 next to the node), we flag the sample as “unstable downstream”.

Built-in Y-SNP trees (as of November 2025) include ISOGG (International Society for Genetic Genealogy) (with 15,786 haplogroups and 65,652 SNPs), YFull (with 40,330 haplogroups and 257,950 SNPs) and FamilyTree DNA (with 96,138 haplogroups and 476,955 SNPs). All SNPs in the tree were filtered to lie in the ∼10 Mb long single-copy, short-read mappable region of the Y-chromosome^19^ and to occur uniquely in the tree topology. GRCh37 support is achieved via liftover of GRCh38 coordinates from the Y-SNP trees using CrossMap^20^. Genotype calling is performed with BCFtools^21^ (mpileup options -B -q 30 -f reference -R chromosome; call options -Am -Oz –-ploidy 1).

### 2.2. Tree-based scoring and heuristic steps

*ybyra* uses a tree-based parsimonious scoring algorithm to assign haplogroups. For each sample, all observed derived and ancestral SNPs are mapped to the selected Y-SNP tree. For each node where either call is made, the pipeline calculates a score, where each derived allele in the path root-to-tip contributes +1 and each ancestral allele -1. The node with the highest cumulative score is selected as the most likely haplogroup, which is followed by a series of heuristic checks to refine the assignment.

When multiple nodes share the highest score, *ybyra* assigns their most recent common ancestor with a derived hit with the new high score. A conservative heuristic that we call the “five-step rule” further requires that the top-scoring node is supported by at least one other derived SNPs within five upstream nodes. If the final assigned node includes an ancestral SNP, the call is flagged in the output as a likely unstable downstream topology.

All intermediate scores, tie resolutions and flags are reported to facilitate manual review as needed, and reproducibility. In addition, plots of both the final haplogroup calls and any scoring ties allow users to quickly inspect results.

### 2.3. Damage filter

On top of the low depth of coverage typical of ancient DNA datasets, ancient DNA has a characteristic substitution pattern deriving from *postmortem* cytosine deamination. This damage profile manifests itself as elevated rates of C→T and G→A transitions at the read termini, depending on library chemistry^22,23^.

*ybyra* includes an optional ancient DNA filter that screens for those substitutions and excludes them during genotype calling step. Separate profiles are implemented for single-stranded and double-stranded library preparations, allowing users to tailor the filter to their library build.

### 2.4 Plotting

All results are printed in tables to allow further fine-grained analyses. Detailed plots of the placement of each sample are produced using ETE3^24^. These allow users to inspect each sample in the context of the relevant branches of the tree. Lastly, *ybyra* includes a dedicated plotting tools called *ybyra-puera*, which uses *genesis*^25^ to produce publication-quality figures from the output of the *ybyra* workflow, visualizing each sample and a summary of all samples on the tree, as in Figure 1.

## 3. Comparison with existing tools

### 3.1. Performance

To evaluate the performance of *ybyra*, we compared it with two other automated haplogroup callers (Yleaf^14^ and haploGrouper^15^) using a dataset consisting of 78 ancient male individuals from the Italian Peninsula^26^.

As input for *ybyra*, we used the FamilyTree DNA Y-SNP tree, which contains almost twice as many SNPs as that used by Yleaf and more than seven time the number used in haploGrouper. Despite the larger number of SNPs analyzed, we found *ybyra* to be substantially faster than both programs – 22% faster than haploGrouper and 45% than Yleaf (**Supplemental Table S1**). It should also be noted that haploGrouper uses VCF files as an input, meaning the most computationally demanding step of the workflow, calling genotypes from the BAM files, is not included in its time estimate.

### 3.2. Concordance and accuracy

We classified haplogroup calls as ‘concordant’ when identical, ancestral or descendant between tools, and ‘discordant’ when on unrelated branches. Calls were 98.1% concordant between *ybyra* and haploGrouper and 92.3% between Yleaf and *ybyra*. All discordant cases corresponded to nodes lacking any derived or ancestral calls in *ybyra*, indicating their placement by Yleaf and haploGrouper relies only on the non-unique recurring SNPs or low-quality markers outside the mappable region excluded by *ybyra*.

Among concordant calls, 57.1% (haploGrouper) and 61.1% (Yleaf) were identical to *ybyra* results. In the remaining cases, the other tools typically placed samples downstream of *ybyra’*s node, despite those having fewer supporting derived SNPs. This confirms that *ybyra*’s heuristics yield the most parsimonious placement (**Supplementary Tables S2-S3**).

### 3.3. Coverage robustness

Finally, we wanted to assess the robustness of *ybyra* to different sequencing depths. We downsampled five high-coverage ancient male genomes^27,28^ to Y-chromosome coverages of 10x, 5x, 2x, 1x, 0.5x, 0.1x, 0.05x and 0.01x, and re-assigned haplogroups using Yleaf and *ybyra*. In all downsampling comparisons, we find haplogroup assignments to be concordant (either identical, ancestral or derived) with the highest coverage assignment (**Supplemental Figure S1, Supplemental Table S4-S5**), a single exception being a distantly related haplogroup being called for one sample in 0.05x coverage by Yleaf. At almost all downsampled coverages in which *ybyra* and Yleaf assign a different haplogroup, we find the *ybyra* assignment to be closer to the highest coverage assignment, the only exception being one case where *ybyra* conservatively assigns one step upstream due to a score tie. We find the *ybyra* results to be robust at coverages as low as 0.05x (corresponding to ∼0.1x nuclear coverage) with four of the five individuals retaining the same or only slightly upstream haplogroup assignments (≤2 nodes upstream). We thus demonstrate that *ybyra* maintains a more stable and reliable classification even under low coverages typical of ancient DNA data.

## 4. Custom Y-chromosome trees

To demonstrate flexibility beyond human data, we applied *ybyra* to the curated horse Y-SNP tree from Bozlak et al.^8^. After reformatting the tree into *ybyra*’s input structure (SNP list, node parent-child relationships and genome coordinates), we reproduced the haplogroup classification for 171 diverse modern horse samples^8,29,30^ reported in the original study^8^ (**Supplemental Table S6**). Applying *ybyra* to two ancient post-Colonial North American horses^7^ yielded haplogroup assignments consistent with reported nuclear results, and further identifies one individual within a lineage specifically associated with Spanish breeds^31^.

## 5. Conclusion

*ybyra* provides a reproducible, transparent and scalable workflow for Y-chromosome haplogroup inference. Its tree-based scoring algorithm and heuristics can be easily integrated to any curated Y-SNP tree topology and are suitable for both modern and ancient DNA data. Its comprehensive outputs and visualization tools facilitate consistent reporting and robust, interpretable haplogroup assignments across diverse datasets.

**Supplementary Figure S1:**
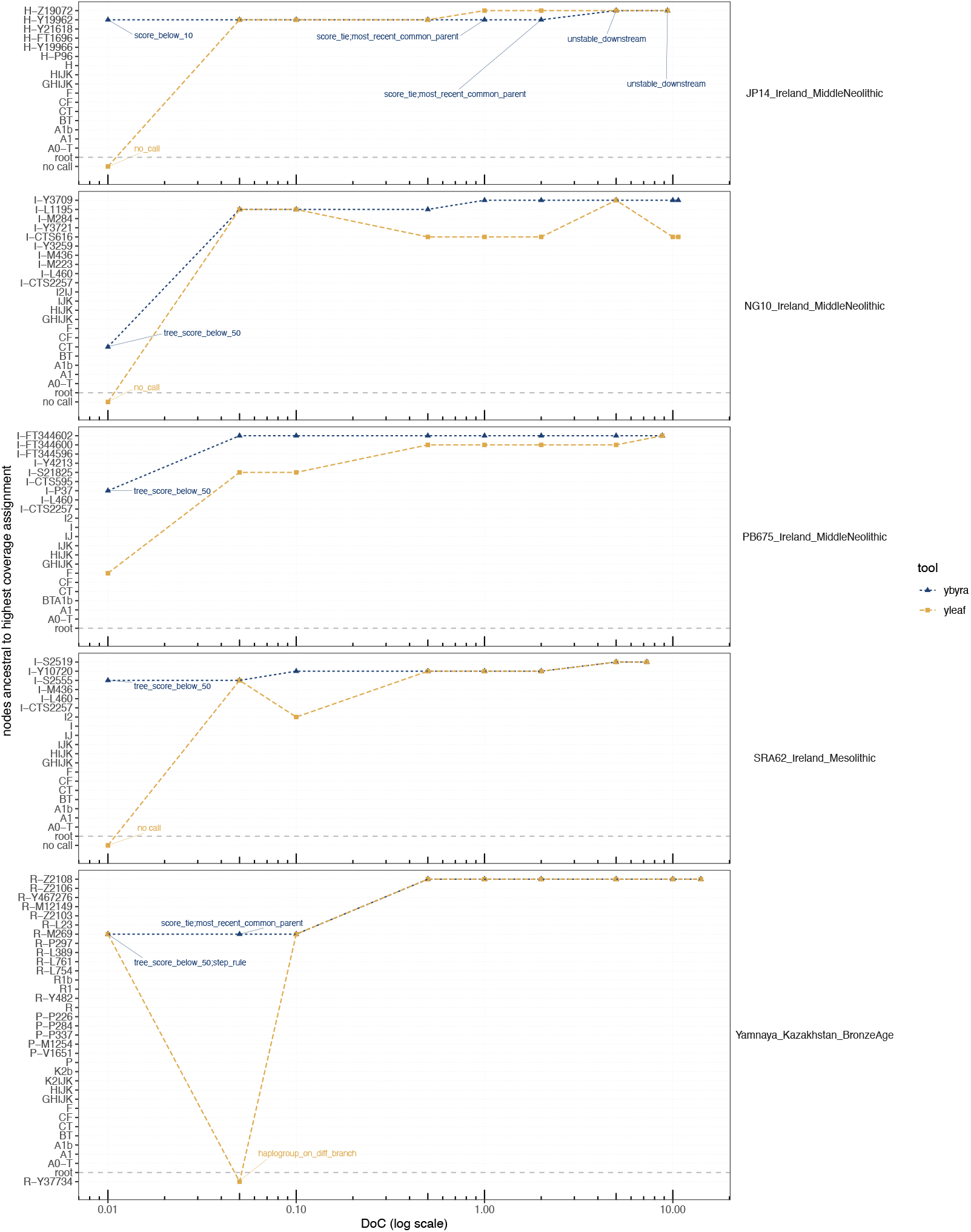
Downsampling comparison experiment. We downsampled 5 high-coverage males to Y-chromosome depth of coverage of 10.0x, 5.0x, 2.0x, 1.0x, 0.5x, 0.1x, 0.05x and 0.01x and used *ybyra* and Yleaf to assign haplogroups. We show that *ybyra* is more robust and stable than Yleaf at all different coverages. The only exception is for JP14_Ireland_MiddleNeolithic, where due to a score tie, *ybyra* conservatively assigns one step upstream.

## Acknowledgements

The authors thank J. Víctor Moreno-Mayar for helpful suggestions, and the ISOGG, YFull and FamilyTree DNA teams for their work curating the human Y-SNP tree and making it publicly available for the research community. The Lundbeck Foundation GeoGenetics Centre is supported by the Lundbeck Foundation (grant nos. R302-2018-2155, R155-2013-16338), the Novo Nordisk Foundation (grant no. NNF18SA0035006), the Wellcome Trust (grant no. UNS69906), Carlsberg Foundation (grant no. CF18-0024), the Danish National Research Foundation (grant nos. DNRF94, DNRF174), the University of Copenhagen (KU2016 programme). All authors were supported by the Novo Nordisk Foundation and Wellcome Trust (AEGIS project) and the Danish National Research Foundation Center for Ancient Environmental Genomics (CAEG).

## References

1. Cortez, D. et al. Origins and functional evolution of Y chromosomes across mammals. Nature 508, 488–493 (2014).

2. Hughes, J. F. & Page, D. C. The Biology and Evolution of Mammalian Y Chromosomes. Annual Review of Genetics 49, 507–527 (2015).

3. Karmin, M. et al. A recent bottleneck of Y chromosome diversity coincides with a global change in culture. Genome Res 25, 459–466 (2015).

4. Poznik, G. D. et al. Punctuated bursts in human male demography inferred from 1,244 worldwide Y-chromosome sequences. Nat Genet 48, 593–599 (2016).

5. Jobling, M. A. & Tyler-Smith, C. Human Y-chromosome variation in the genome-sequencing era. Nat Rev Genet 18, 485–497 (2017).

6. Petr, M. et al. The evolutionary history of Neanderthal and Denisovan Y chromosomes. Science 369, 1653–1656 (2020).

7. Taylor, W. T. T. et al. Early dispersal of domestic horses into the Great Plains and northern Rockies. Science 379, 1316–1323 (2023).

8. Bozlak, E. et al. Refining the evolutionary tree of the horse Y chromosome. Sci Rep 13, 8954 (2023).

9. Underhill, P. A. et al. Y chromosome sequence variation and the history of human populations. Nat Genet 26, 358–361 (2000).

10. Y-Chromosome Consortium. A Nomenclature System for the Tree of Human Y-Chromosomal Binary Haplogroups. Genome Res. 12, 339–348 (2002).

11. Fowler, C. et al. A high-resolution picture of kinship practices in an Early Neolithic tomb. Nature 1–4 (2021) doi:10.1038/s41586-021-04241-4.

12. Seersholm, F. V. et al. Repeated plague infections across six generations of Neolithic Farmers. Nature 632, 114–121 (2024).

13. Poznik, G. D. Identifying Y-chromosome haplogroups in arbitrarily large samples of sequenced or genotyped men. 088716 Preprint at 10.1101/088716 (2016).

14. Ralf, A., Montiel González, D., Zhong, K. & Kayser, M. Yleaf: Software for Human Y-Chromosomal Haplogroup Inference from Next-Generation Sequencing Data. Mol Biol Evol 35, 1291–1294 (2018).

15. Jagadeesan, A. et al. HaploGrouper: a generalized approach to haplogroup classification. Bioinformatics 37, 570–572 (2021).

16. Martiniano, R., De Sanctis, B., Hallast, P. & Durbin, R. Placing Ancient DNA Sequences into Reference Phylogenies. Mol Biol Evol 39, msac017 (2022).

17. Köster, J. & Rahmann, S. Snakemake—a scalable bioinformatics workflow engine. Bioinformatics 28, 2520–2522 (2012).

18. Li, H. et al. The Sequence Alignment/Map format and SAMtools. Bioinformatics 25, 2078–2079 (2009).

19. Poznik, G. D. et al. Sequencing Y Chromosomes Resolves Discrepancy in Time to Common Ancestor of Males versus Females. Science 341, 562–565 (2013).

20. Zhao, H. et al. CrossMap: a versatile tool for coordinate conversion between genome assemblies. Bioinformatics 30, 1006–1007 (2014).

21. Danecek, P. et al. The variant call format and VCFtools. Bioinformatics 27, 2156–2158 (2011).

22. Briggs, A. W. et al. Patterns of damage in genomic DNA sequences from a Neandertal. PNAS 104, 14616–14621 (2007).

23. Gansauge, M.-T. & Meyer, M. Single-stranded DNA library preparation for the sequencing of ancient or damaged DNA. Nat Protoc 8, 737–748 (2013).

24. Huerta-Cepas, J., Serra, F. & Bork, P. ETE 3: Reconstruction, Analysis, and Visualization of Phylogenomic Data. Mol Biol Evol 33, 1635–1638 (2016).

25. Czech, L., Barbera, P. & Stamatakis, A. Genesis and Gappa: processing, analyzing and visualizing phylogenetic (placement) data. Bioinformatics 36, 3263–3265 (2020).

26. Antonio, M. L. et al. Ancient Rome: A genetic crossroads of Europe and the Mediterranean. Science 366, 708–714 (2019).

27. de Barros Damgaard, P. et al. The first horse herders and the impact of early Bronze Age steppe expansions into Asia. Science 360, eaar7711 (2018).

28. Cassidy, L. M. et al. A dynastic elite in monumental Neolithic society. Nature 582, 384– 388 (2020).

29. Felkel, S. et al. The horse Y chromosome as an informative marker for tracing sire lines. Sci Rep 9, 6095 (2019).

30. Remer, V. et al. Y-Chromosomal Insights into Breeding History and Sire Line Genealogies of Arabian Horses. Genes 13, 229 (2022).

31. Radovic, L. et al. The global spread of Oriental Horses in the past 1,500 years through the lens of the Y chromosome. Proceedings of the National Academy of Sciences 121, e2414408121 (2024).

